# ATG-9-Induced Lysosomal Membrane Permeabilization and Cell Death in a *Caenorhabditis elegans* model of Mucolipidosis type IV

**DOI:** 10.64898/2026.07.06.736802

**Authors:** Hope Dang, Teresa Horm, Savannah Perno, Shirine Gholam, Marvin O’Ketch, Shaharyar Ashraf, Sebastian Hernandez, Justin Randall, Hanna Fares

## Abstract

Mucolipidosis type IV is a lysosomal storage disease that is characterized by delayed psychomotor development and retinal degeneration due to cell death, in addition to other symptoms that are due to aberrant functions of live tissues. *Caenorhabditis elegans* CUP-5 is the orthologue of human TRPML1, the protein that is dysfunctional in Mucolipidosis type IV patients. Mirroring Mucolipidosis type IV pathology, loss of *C. elegans* CUP-5 results in developing intestinal cell death in embryos leading to embryonic lethality, while other tissues in adults lacking CUP-5 are alive but dysfunctional. We had previously shown that ESCRT-Associated proteins and the ATP-Binding Cassette Transporter MRP-4 are necessary for acquiring aberrant and poorly functional lysosomes in the absence of CUP-5. In this study, we show that the aberrant lysosomes permeabilize or rupture, thus releasing lysosomal degradative enzymes that kill cells in the absence of CUP-5. We also show that the autophagy-related protein ATG-9 mediates, in an autophagy-independent manner, this lysosomal permeabilization. We finally propose phenotypic and biochemical models linking CUP-5 to lysosomal defects and cell death.

## Introduction

Mucolipidosis type IV (MLIV) is a lysosomal storage disorder that is caused by mutations in the gene *MCOLN1* that encodes the cation channel protein TRPML1/mucolipin-1 [1–3]. Almost all tissues in MLIV patients have enlarged “lysosomes” [4–7], these giant “lysosomes”, referred to henceforth as lysosomes, are enlarged hybrid organelles that have components of at least lysosomes, late endosomes, and autophagosomes [8–10]. MLIV patients present with two classes of symptoms. First, progressive and developmental brain neuronal and retinal degeneration that lead to psychomotor impairment and problems with vision, respectively [11–13]. Indeed, siRNA-mediated loss of TRPML1 in tissue-culture cells results in the specific leakage of Cathepsin B from lysosomes leading to the activation of apoptosis [14]. Second, defective functions of otherwise living tissues that lead to symptoms like corneal clouding and achlorhydria [15].

CUP-5 is the *Caenorhabditis elegans* orthologue of TRPML1. Mutations in *cup-5* result in large abnormal lysosomes in most *C. elegans* tissues [16, 17]. Similar to MLIV patients, *cup-5* mutants have two classes of phenotypes. First, embryos laid by *cup-5(null)* hermaphrodites fail to hatch due to the death of developing intestinal cells [17–19]. Second, adult tissues do not undergo death/degeneration even though they have abnormal membrane transport [8, 20]. In the absence of CUP-5, the lysosomal defect in developing intestinal cells is molecularly unrelated to the one in adult tissues because genetic suppressors of the developing intestinal cell lysosomal phenotype do not suppress adult tissue lysosomal defects [20]. Expression of human TRPML1 in worms lacking CUP-5 suppresses all of the phenotypes, indicating that *C. elegans cup-5* is a good model of MLIV [8, 18].

What is the basis for the neuronal death in the absence of TRPML1? A variety of cellular defects have been shown in the absence of TRPML1, including defects in heavy metal homeostasis, mitochondrial aberrations, production of reactive oxygen species, and dysfunction in different aspects of membrane transport/organelle biogenesis [9, 21–34]; yet, it is not clear which of these defects contribute to neural degeneration. As mentioned above, siRNA-mediated knockdown of TRPML1 in HeLa cells results in Cathepsin B leakage from lysosomes and Bax-mediated apoptosis [14]. In contrast, in *C. elegans* lacking CUP-5, embryonic viability is only partially suppressed by blocking caspase-mediated apoptosis [17, 18]. Indeed, we showed that loss of CED-3, or increasing ATP levels in embryos lacking CUP-5, does not suppress the lysosomal defects and only results in a mild increase in embryos hatching which subsequently arrest at the L1 larval stage [17]. This suggested that *in vivo* in worms, conventional apoptosis is not the major basis for the developing intestinal cell death.

Genetic suppression is a powerful tool to uncover pathways leading to cell death in the absence of CUP-5. We identified two classes of extragenic suppressors of the lysosomal defects and the developing intestinal cell death, ESCRT-Associated proteins and the ATP-Binding Cassette (ABC) Transporter MRP-4 [19, 20]. RNAi knockdown or null alleles of the encoding genes almost fully suppress the developing intestinal cells death, and the embryonic and larval viability, of worms lacking CUP-5. This indicates that ESCRT-Associated proteins and MRP-4 are necessary for the lysosomal defects and the death of developing intestinal cells in the absence of CUP-5. Our results are consistent with a model where, in the absence of CUP-5, there is an increased activity of the ESCRT-Associated de-ubiquitinase protein USP-50 (homologue of human USP8/UBPY) leading to the hypo-ubiquitination, and hence increased activity, of MRP-4 on endosomes/lysosomes: this MRP-4 hyper-activation causes the lysosomal defect in developing intestinal cells [19, 20]. Once there is a lysosomal defect in the absence of CUP-5, what subsequently causes the death/degeneration?

In this study, we describe another suppressor, *atg-9*, of the *cup-5* mutant developing intestinal cell death: ATG-9 functions downstream of the lysosomal defect. Our studies suggest that in the absence of CUP-5, the lysosomal defect triggers an ATG-9-mediated lysosomal permeabilization/rupture that causes the degeneration of developing intestinal cells.

## Materials and methods

### *C. elegans* strains and methods

Standard methods were used for the growth and manipulation of *C. elegans,* var. “Bristol” strain [35]. *C. elegans*, grown on NGM plates (with OP50), was manipulated using a dissection microscope sometimes equipped with fluorescence optics (Zeiss, Thornwood, NY) [35]. The wild type var. “Hawaiian” strain was used for single nucleotide polymorphism mapping, as previously described [36]. RNAi was done by the feeding method; control RNAi was done using bacteria carrying the double-stranded RNA generating vector L4440/pPD129.36 [37]. Two *cup-5* null alleles were used: *n3194* and *zu223* [18]. The *C. elegans* strains used in this study are listed in Table 1. The following transgenes were used in this study:

*- arIs37[*P*myo-3::ssGFP; dpy-20]*: expresses GFP that is secreted by body wall muscles into the body cavity and is endocytosed primarily by coelomocytes; *cup-5* mutants result in fluorescently brighter coelomocytes due to a defect in the degradation of GFP in lysosomes of coelomocytes [38].
*- bIs1[vit-2::GFP; rol-6(su1006)]*: expresses the yolk protein VIT-2::GFP fusion that is secreted by adult intestinal cells and is endocytosed primarily by oocytes [39].
*- pwIs50[lmp-1::GFP, unc-119]:* expresses LMP-1, the worm homologue of mammalian Lamp1, fused to GFP [8].
*- kxEx141(cpr-6::mCherry; rol-6(su1006)]:* expresses CPR-6, the worm homologue of mammalian Cathepsin B, fused to mCherry [40].
*- kxEx148(gba-3::mCherry; rol-6(su1006)]:* expresses GBA-3, the worm homologue of mammalian Glucosylcerebrosidase, fused to mCherry [40].
*- cdIs260[*P*elt-2::MRP2(human);* P*myo-2::GFP; unc-119]*: was made using ballistic transformation and expresses the human ABC Transporter MRP2 under the control of the intestine-specific *elt-2* promoter [41, 42]. The plasmid encoding MRP-2 under the control of the *elt-2* promoter is pHD845; details of construction are available upon request.

**Table 1.**
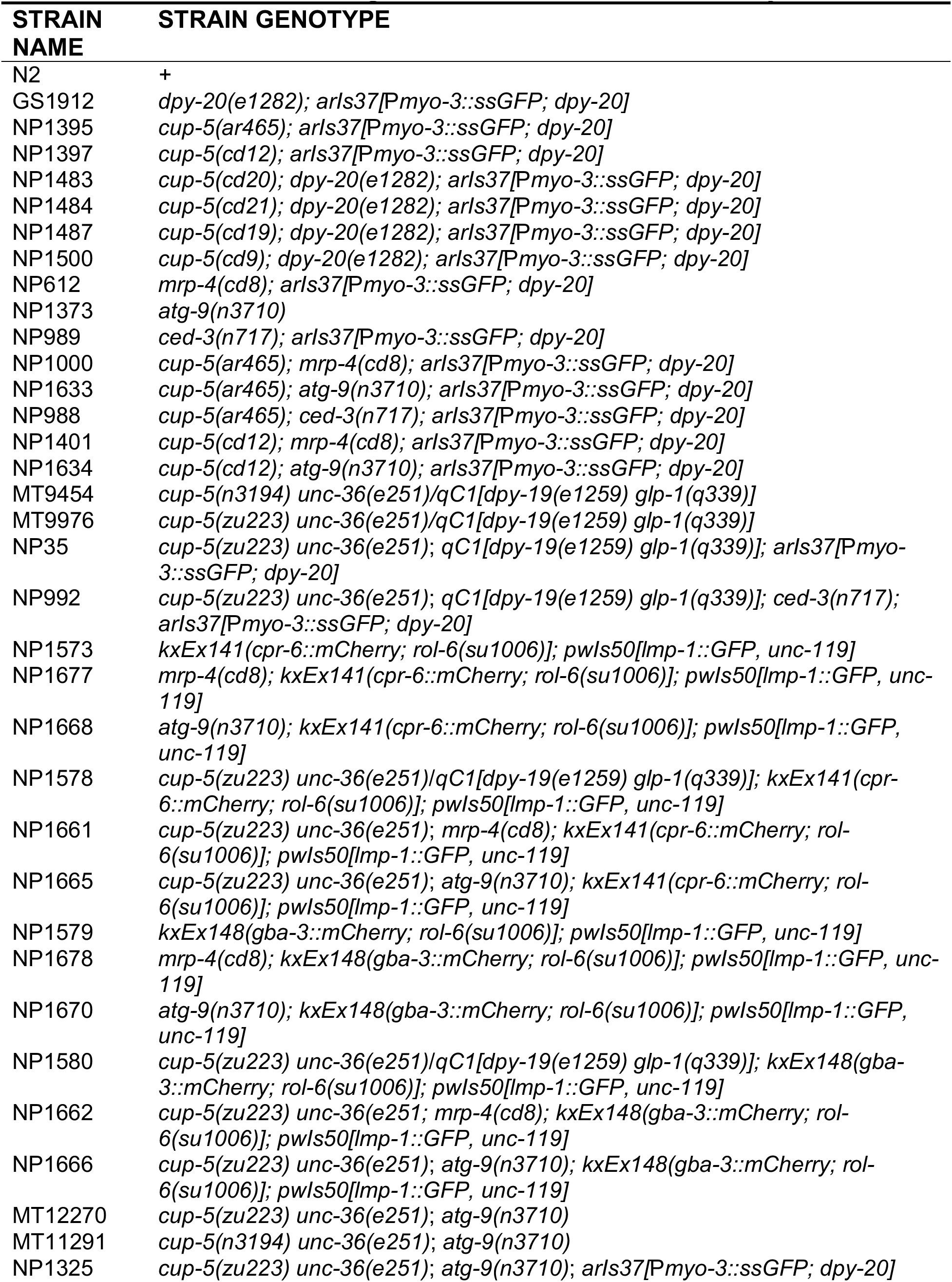
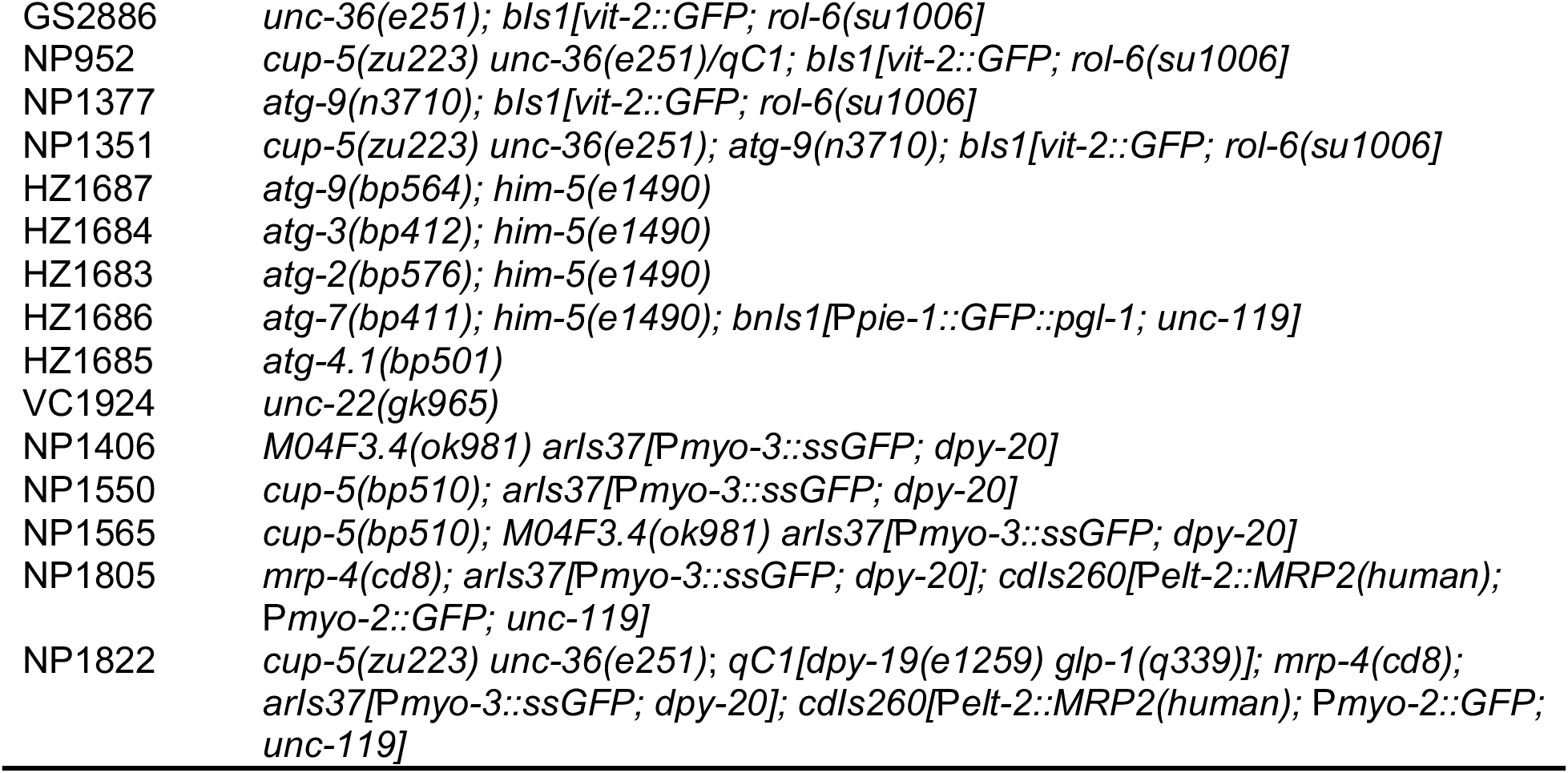
*C. elegans* strains used in this study.

### Measurement of embryonic viability

Each viability measurement consists of at least three independent experiments for statistical analysis. The results show the averages and standard deviations.

#### Normal growth

Adult worms were allowed to lay eggs overnight on NGM + OP50 plates at 20 °C. The adults were then removed and the percentage of eggs that hatched and developed to adults was determined.

#### RNA interference (RNAi)

For RNAi conditions, embryonic viability was measured on NGM + Carbenicillin (50 µg/ml) + IPTG (1 mM) plates seeded with bacteria expressing the desired double-stranded RNA. In addition, adult worms were moved after one day to a second plate with the same bacterium and embryonic viability was determined from the second plate (adult was removed after one day of laying eggs).

#### MSDH + RNAi

*O*-methyl-serine dodecylamide hydrochloride (MSDH) was dissolved in DMSO at a concentration of 50 mM. First, we determined the highest concentration of DMSO that does not affect the viability of wild type worms, using plates with the same amounts of DMSO as a control. The MDSH + RNAi experiments were done using a similar procedure as RNAi experiments, except that MSDH was added to the experimental plates, when the medium had cooled down before pouring the plates, at a final concentration of 50 µM; plates without MSDH had the same amount of DMSO included in the medium.

### Identification of *n3710*, the *cup-5* suppressor mutation

Ethyl methane sulfonate (EMS) mutagenesis was done on strain MT9454: *cup-5(n3194) unc-36(e251)/qC1[dpy-19(e1259) glp-1(q339)]* as previously described [18, 19]. Briefly, *cup-5(n3194)* is a maternal effect embryonic lethal such that homozygous *cup-5(n3194)* hermaphrodites lay almost 100% embryos that do not hatch [18]; *unc-36(e251)* yields uncoordinated movement and is closely linked to *cup-5* [35]; *qC1[dpy-19(e1259) glp-1(q339)]* is a balancer chromosome that suppresses recombination between *cup-5* and *unc-36* [43].

Normally-moving MT9454 worms were EMS-mutagenized and were allowed to self-fertilize. F_2_ Unc hermaphrodites were then placed on separate plates; viable F_3_ progeny indicates the presence of an intragenic or extragenic suppressor, distinguished by crossing to *cup-5(n3194) unc-36(e251)/qC1[dpy-19(e1259) glp-1(q339)]* males and examining the F_2_ progeny of F_1_ Unc cross-progeny: intragenic suppressors result in viable F_2_ progeny while recessive extragenic suppressors result in F_2_ embryos that fail to hatch. The *cup-5* extragenic suppressor *n3710* was identified from such a pilot screen.

### *atg-9* complementation analysis

Male HZ1687: *atg-9(bp564); him-5(e1490)* or wild type N2 worms were crossed to NP621: *cup-5(zu223) unc-36(e251); arIs37[*P*myo-3::ssGFP; dpy-20]; n3710* hermaphrodites (Fig 1A).

**Fig 1.**
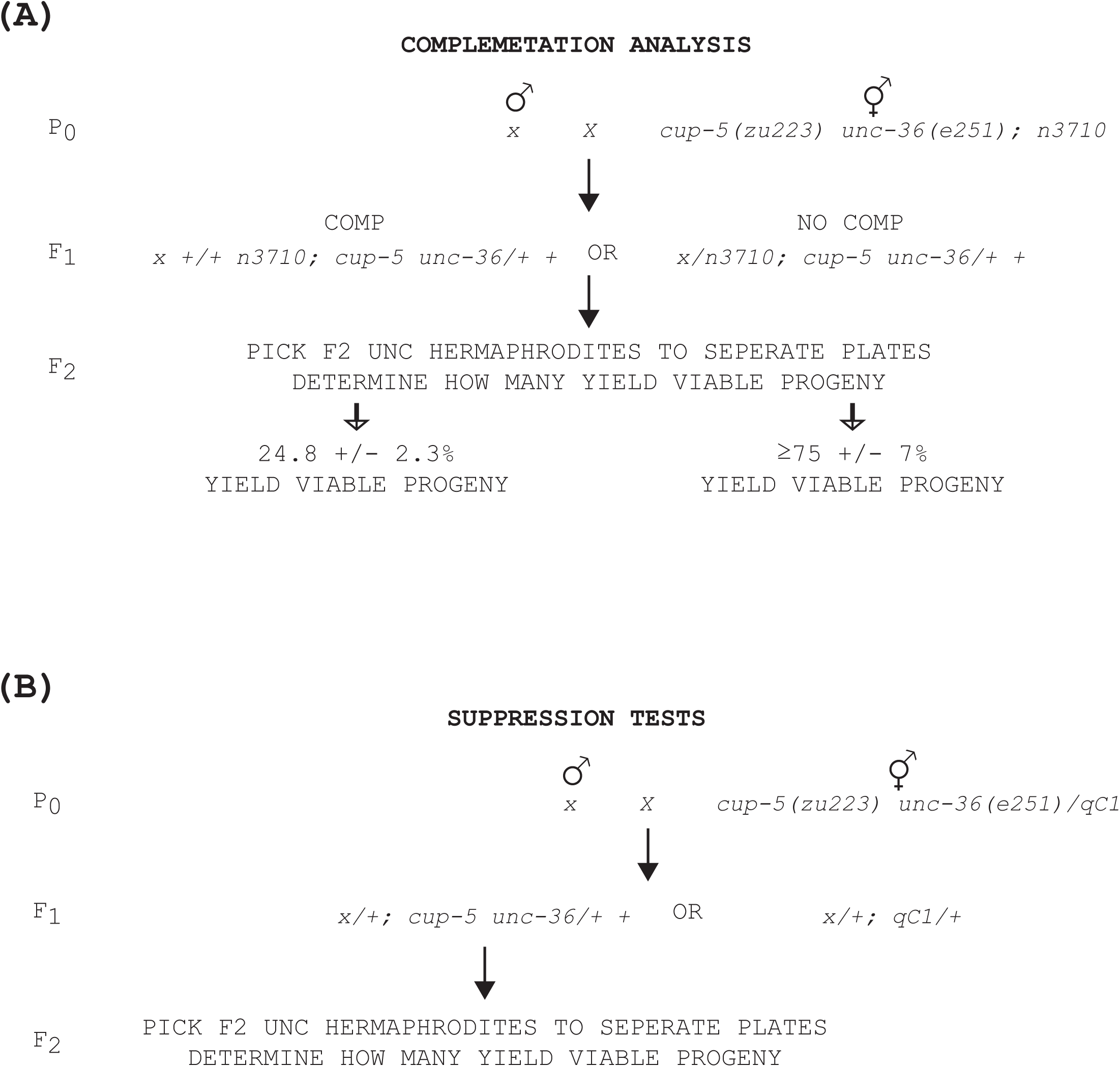
Schematic of complementation and suppression tests. (A) Complementation analysis testing whether *n3710* corresponds to a mutation in *atg-9*. “*x*” in P0 represents wild type or *atg-9(bp564)*. The predicted percent of F2 that lay viable progeny is adjusted for the percent *n3710* suppression of *cup-5(null)* lethality. (B) Cross used to determine whether mutations are able to suppress the embryonic lethality caused by the *cup-5(zu223)* null allele. “*x*” in P0 represents wild type or various mutations.

Independent *cup-5 unc-36* F_2_ hermaphrodites were picked to separate NGM + OP50 plates and the percentage of hermaphrodites that yielded viable progeny was determined (Fig 1A). The percentage F_2_ parents that lay viable is theoretically ∼33% for complementation and 100% for non-complementation. However, *n3710; cup-5(null)* lay 75 +/- 7% viable progeny, so the prediction for complementation is 24.8 +/- 2.3% and for non-complementation is at least (depending on the strength of *bp564* suppression of *cup-5* lethality) 75 +/- 7%.

### cup-5(zu223) suppression tests

Males of wild type N2 or strains bearing mutations of interest were crossed to NP35: *cup-5(n3194) unc-36(e251)/ qC1[dpy-19(e1259) glp-1(q339)]; arIs37[*P*myo-3::ssGFP; dpy-20]* hermaphrodites (Fig 1B). At least 100 independent *cup-5 unc-36* F_2_ hermaphrodites were placed on separate NGM + OP50 plates and the percentage of F_2_ hermaphrodites that yielded viable progeny was determined (Fig 1A). For each mutant strain, the cross was repeated independently three times for statistical analysis. The results show the averages and standard deviations.

### Embryo immunofluorescence

Immunofluorescence staining to detect the yolk receptor RME-2, the Notch Receptor LIN-12, and the intermediate filament protein IFB-2 in embryos was done as previously described [44].

### Microscopy

Worms were placed in 6 μl of 9 mM levamisole/1X PBS on a 2.2% agarose pad. Confocal images were taken on a Zeiss LSM510 (Carl Zeiss, Oberkochen, Germany) laser scanning confocal, using LSM imaging software. Image analysis was done using Metamorph software (Molecular Devices, Sunnyvale, CA). For VIT-2::GFP, RME-2, and LIN-12, wild type and mutant strain images were taken in the same session using identical microscopy settings to allow for comparison of fluorescence signal strengths. For VIT-2::GFP, at least 12 embryos for each strain were imaged and all of the VIT-2::GFP compartments within developing intestinal cells of each embryo were measured to determine averages and standard deviations.

### Organelle fractionation of embryos

For each strain, two plates of NGM + OP50, each containing 1000 adult hermaphrodites that were rollers and/or showed red fluorescence (indicating the presence of *kxEx* transgenes) were left at 20 °C for one day to lay eggs; Unc worms, thus homozygous for *cup-5*, were chosen from *cup-5 unc-36/qC1* strains. Parallel wild type and *cup-5(zu223)* mutant strains carrying the same *kxEx* transgene were processed at the same time to ensure identical treatments, especially sonication to disrupt embryos and cells. The next day, the worms and eggs were collected and were treated twice with bleach solution (1% NaClO, 1 M NaOH), each time for 5 minutes, to dissolve the OP50 and adult worms. The remaining embryos were washed three times in M9 buffer (22 mM KH_2_PO_4_, 43 mM Na_2_HPO_4_, 86 mM NaCl, and 1 mM MgSO_4_) and were transferred to a 1.5 ml Eppendorf (Hamburg, Germany) tube. The embryos were microcentrifuged at 3000*g*, the supernatant was removed, and the embryos were resuspended in 0.3 ml of ice-cold Membrane Lysis Buffer (0.2 M Sorbitol, 50 mM KCH_3_COO, 2 mM EDTA, 2 mM HEPES-KOH, 1 mM DTT, pH 6.8; right before use, 1 tablet of Roche Life Sciences [Indianapolis, IN] Complete Protease Inhibitor and 20 μl of 1 M N-Ethylmaleimide were added per 1 ml of buffer). The eggs were immediately sonicated using a Bioruptor (Diagenode, Denville, NJ) sonicator for 1 second at high power followed by 5 seconds at low power. The tubes were then centrifuged at 3000*g* for 15 minutes at 4 °C and the supernatants were transferred to separate tubes (the pellets represent large debris like embryos that weren’t fully disrupted). The supernatants were microcentrifuged at 200,000*g* for 1 hour at 4 °C. The pellets from this spin, that represent the membrane fractions, were resuspended in 200 μl of 1X Laemmli Buffer-Urea (4% SDS, 1% NP-40, 5% β-mercaptoethanol, 3 M Urea, 10 mM DTT, 10% glycerol, 0.002% bromophenol blue, 0.05 M Tris-HCl, pH 6.8; right before use, 1 tablet of Roche Life Sciences Complete Protease Inhibitor was added per 1 ml of buffer). The supernatants from the 200,000*g* spin were concentrated using Pierce Protein Concentrators (ThermoFisher Scientific, Rockford, IL) to a volume of 100 μl and 100 μl of 2X Laemmli Buffer-Urea was added to make a final volume of 200 μl (same as the pellet). Samples were analyzed on Western Blots. The intensities of the bands were determined using ImageJ (JACoP; National Institute of Health, Bethesda, MD) [45].

### Molecular methods

Standard methods were used for the manipulation of recombinant DNA [46]. Polymerase chain reaction was done using the Expand Long Template PCR System (Roche Life Sciences) according to the manufacturer’s instructions. All other enzymes were from New England Biolabs (Beverly, MA).

### Whole-genome sequencing

Strain NP1325: *cup-5(zu223) unc-36(e251)*; *n3710*; *arIs37[*P*myo-3::ssGFP; dpy-20]* was made by backcrossing strain MT12270: *cup-5(zu223) unc-36(e251)*; *n3710* ten times to the wild type GS1912: dpy-20(e1282); *arIs37[*P*myo-3::ssGFP; dpy-20]* strain. NP1325 was then crossed one more time to GS1912 (*dpy-20(e1282); arIs37[*P*myo-3::ssGFP; dpy-20]*) and ten independently isolated homozygous *cup-5(zu223) unc-36(e251)*; *n3710*; *arIs37[*P*myo-3::ssGFP; dpy-20]* hermaphrodites were picked from the F_3_ generation. These ten isolates were mixed together for genomic DNA isolation for sequencing; this allowed us to identify background mutations that would not be homozygous in all ten isolates (unless they are closely linked to the *cup-5, unc-36, n3710, or arIs37*). The genome of strain MT11291: *cup-5(n3194) unc-36(e251)*; *n3710* was also sequenced. Mutations that were found in both NP1325 and MT11291 (besides *cup-5* and *unc-36*) but that were absent in GS1912 and the *C. elegans* reference genome were considered as candidate *n3710* mutations. *C. elegans* genomic DNA was sequenced by the University of Arizona Genomics Core using shotgun/de novo sequencing on an Illumina HiSeq 2000. Bowtie2 was used to map reads from each sample to the *C. elegans* reference WBcel215. samtools mpileup was then used to identify all of the variations from WBcel215. The mutant sequence was compared to the whole genome sequence of GS1912; unique variants each had an average of 147 reads. This identified homozygous mutations (including *cup-5* and *unc-36*) relative to the parent strain GS1912.

### Statistical methods

The Student’s *t*-test was used to compare average measurements from two samples using a two-tailed distribution (Tails=2) and a two-sample unequal variance (Type=2).

### Data availability

All data and reagents are available upon request.

## Results and discussion

### Lysosomal rupture in the absence of *cup-5*

We hypothesized that in the absence of CUP-5, the large lysosomes may rupture or leak hydrolytic enzymes that would degrade cellular components leading to tissue degeneration. We therefore determined the localization of two lysosomal enzymes in developing intestinal cells of embryos: CPR-6 (homologue of human Cathepsin B) and GBA-3 (homologue of human Glucosylcerebrosidase) [40]. In wild type developing intestinal cells, both enzymes co-localized with LMP-1, confirming their lysosomal localization (arrows in Fig 2A). In embryos lacking CUP-5, in addition to this co-localization with LMP-1 on lysosomes, some developing intestinal cells also showed diffuse cytoplasmic localization of CPR-6 and GBA-3 (arrowheads in Fig 2A). Thus, it appeared that at least two lysosomal enzymes localize to the cytoplasm of *cup-5(null)* developing intestinal cells.

**Fig 2.**
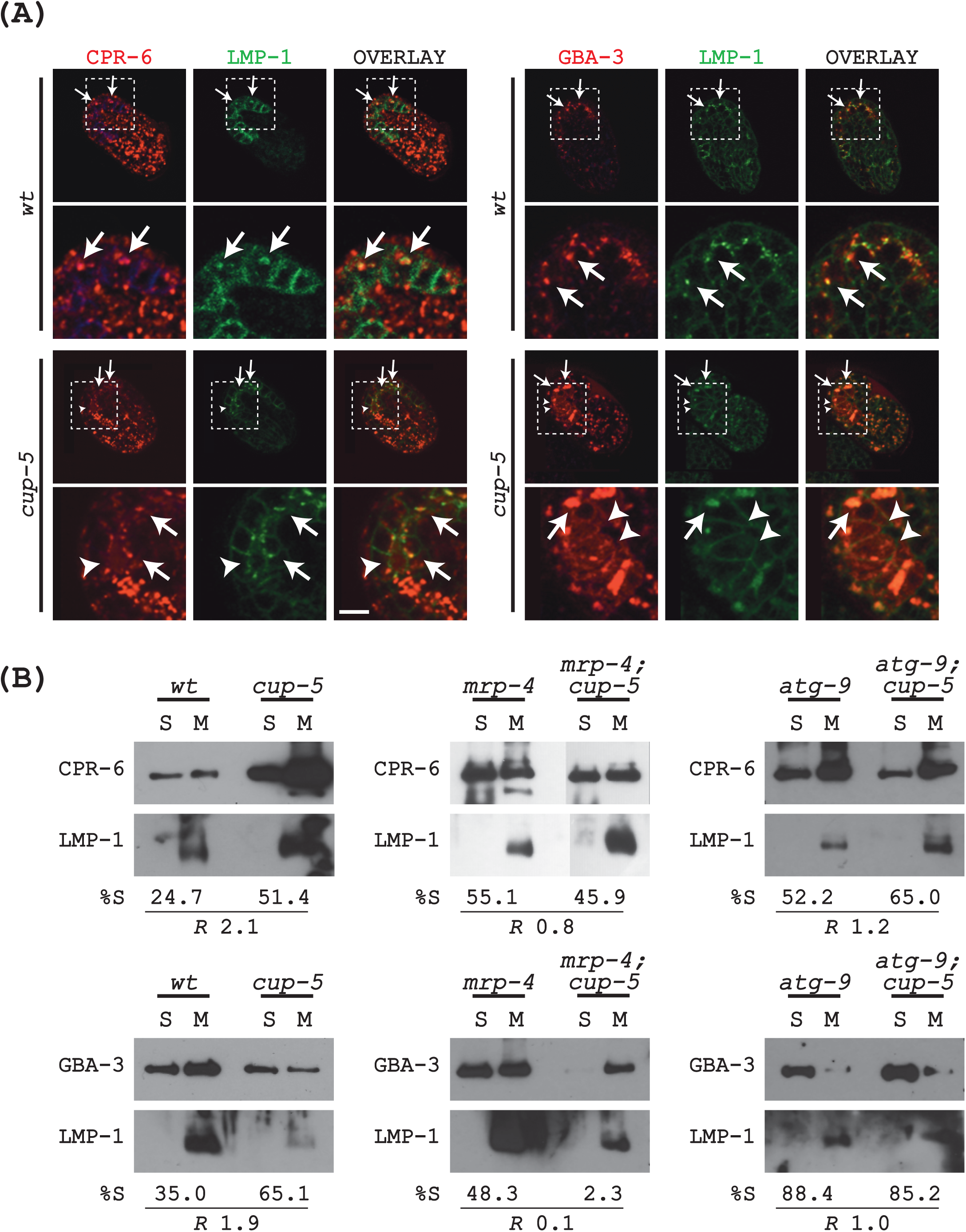
Lysosomal permeabilization in the absence of CUP-5. (A) Confocal images of “comma” to “1.5-fold” stage embryos laid by wild type or *cup-5(null)* hermaphrodites expressing CPR-6::mCherry or GBA-3::mCherry, and LMP-1::GFP. The boxed area is digitally zoomed. Arrows indicate co-localization of mCherry and GFP signals; arrowheads indicate cytoplasmic localization of mCherry. Scale bar represents 10 µm in whole embryos. (B) Western blot following organelle fractionation of embryos of the indicated genotypes expressing CPR-6::mCherry or GBA-3::mCherry, and LMP-1::GFP. S, soluble/cytoplasmic fraction; M, organelle-associated fraction. LMP-1::GFP is an integral membrane protein that we used to ascertain proper fractionation. The percent signal in the soluble fraction relative to the total signal (S + P) is indicated. R represents the ratio of %S from the *cup-5(zu223)* strain divided by the %S from the corresponding “wild type” strain.

We used organelle fractionation of embryos to confirm this cytoplasmic leakage of lysosomal lumenal proteins. The same strains used for imaging were biochemically manipulated to separate organelle-associated (M) from cytoplasmic soluble (S) proteins. Wild type strains showed some CPR-6 and GBA-3 in the cytoplasm (S fraction), likely due to partial rupture during the procedure. The percentage of CPR-6 and GBA-3 in the cytoplasm was two times higher in the absence of CUP-5 than in wild type embryos (Fig 2B). If this lysosomal enzyme leakage is associated with lethality in the absence of CUP-5, then it should be suppressed by eliminating MRP-4. Loss of MRP-4 suppresses the embryonic lethality of *cup-5(null)* hermaphrodites: *mrp-4(null); cup-5(null)* double mutant worms result in ∼90% viable and healthy progeny, as opposed to almost 0% for *cup-5(null)* single mutant worms [19]. Indeed, *mrp-4; cup-*5 embryos do not show increased cytoplasmic fractionation of either CPR-6 or GBA-3 relative to *mrp-4* single mutant embryos (Fig 2B).

Our results are consistent with the results in HeLa cells showing lysosomal leakage after reducing TRPML1 levels [14]. However, while only Cathepsin B leaked out of lysosomes in HeLa cells, our results suggest a more general leakage of lysosomal enzymes into the cytoplasm in the absence of CUP-5 in *C. elegans*. The question became whether this leakage contributes to the death of developing intestinal cells in the absence of CUP-5.

### Physiological significance of lysosomal rupture

*O*-methyl-serine dodecylamide hydrochloride (MSDH) is a lysosomotropic detergent that destabilizes lysosomal membranes leading to the lysosomal leakage [47]. We reasoned that if lysosomal leakage contributed to *cup-5* mutant cell death, then MSDH would exacerbate the cell, and therefore embryonic, death of *cup-5* hypomorphic mutants. We first identified 50 µM as the highest concentration of MSDH that wasn’t lethal to wild type worms; at this concentration, 100% of wild type embryos hatched and grew to fertile adults (Fig 3A). However, MSDH did not significantly increase the embryonic lethality of several independently isolated *cup-5* hypomorphic alleles; these *cup-5* alleles lay 100% viable embryos that grow to adults under normal conditions (Fig 3A) [16, 48].

**Fig 3.**
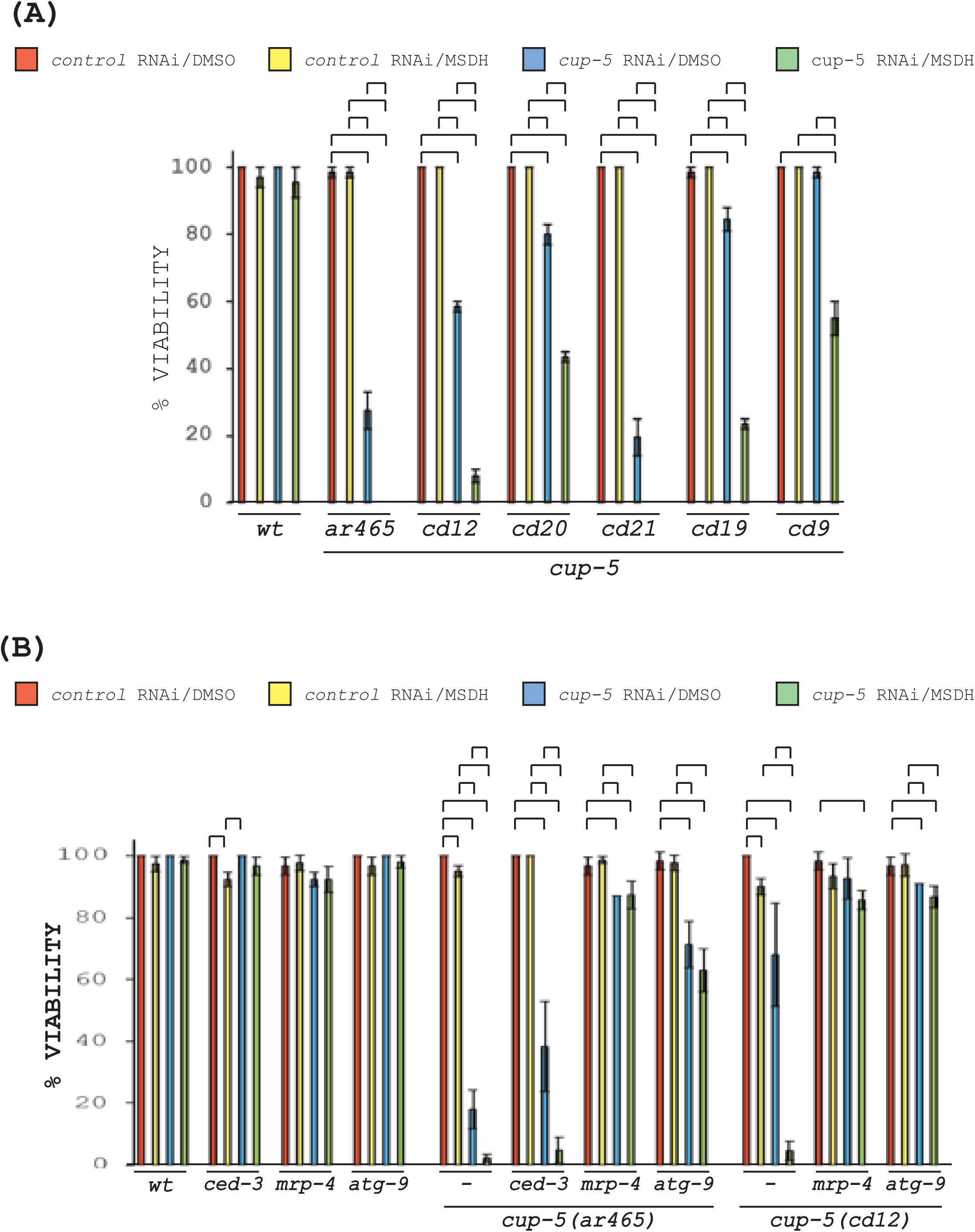
MSDH-induced synthetic lethality in the absence of CUP-5. (A) Percent of viable worms laid by wild type or *cup-5* hypomorphic mutant hermaphrodites after control (L4440) or *cup-5* RNAi and in the presence of MSDH (50 µM) or an equal amount of DMSO. Inverted brackets indicate *P<*0.05. All strains also had the *arIs37* transgene. (B) Percent of viable worms laid by the indicated wild type and mutant hermaphrodites after control (L4440) or *cup-5* RNAi and in the presence of DMSO or MSDH as in “A”. Inverted brackets indicate *P<*0.05. All strains except *atg-9(n3710)* also had the *arIs37* transgene.

We therefore used *cup-5* RNAi to further reduce the levels of CUP-5 in worms. *cup-5* RNAi did not affect the embryonic viability of wild type embryos in the presence or the absence of MSDH (Fig 3A). As expected, *cup-5* RNAi reduced the viability of *cup-5* hypomorphic embryos (except for the weakest allele *cd9*), essentially driving them towards the *cup-5(null)* phenotype of 0% embryonic viability (Fig 3A) [17, 18]. Importantly, this reduction in embryonic viability following *cup-5* RNAi was significantly exacerbated by MSDH in every *cup-5* hypomorphic strain (Fig 3A). Thus, MSDH specifically enhances the embryonic death due to reduced CUP-5 levels.

If this MSDH-induced lethality is physiologically relevant to the embryonic lethality in *cup-5(null)* embryos, then it should be suppressed by extragenic suppressors of *cup-5(null)* embryonic lethality. Indeed, an *mrp-4(null)* mutation eliminated the MSDH-induced lethality of *cup-5(ar465)* and *cup-5(cd12)* hypomorphic alleles following *cup-5* RNAi (Fig 3B). In contrast, *ced-3(null)* mutations only partially suppress *cup-5(null)* lethality such that *ced-3(null); cup-5(null)* double mutant worms result in ∼5% of embryos that hatch but arrest as L1 larvae [17]. Consistent with this result, a *ced-3(null)* did not affect the MSDH-induced lethality of *cup-5(ar465)* following *cup-5* RNAi (Fig 3B). Thus, MSDH exacerbates lethality when CUP-5 levels are reduced by enhancing the same physiological response that leads to cell death in the absence of CUP-5.

These results suggest that the mechanism for cell death in the absence of CUP-5 is the rupture or permeabilization of lysosomes resulting in the leakage of lysosomal enzymes that digest cellular components. Lysosome size is significantly larger and lysosomal functions are compromised in cells that lack CUP-5, which could trigger this permeabilization. The question remained whether this lysosomal defect automatically results in rupture/permeabilization, or whether this rupture is mediated by a protein or pathway.

### Identification of *atg-9*, an extragenic suppressor of *cup-5(null)* lethality

The *n3710* mutation was identified as an extragenic suppressor of *cup-5(null)* embryonic lethality (B. Hersh, and H.R. Horvitz, personal communication). Using single nucleotide polymorphism mapping, we mapped *n3710* to the left of F36H9 on chromosome V, approximately 1700 kb from the left end of chromosome V. We then did whole genome sequencing of two independent strain bearing the *n3710* mutation: besides *cup-5* and *unc-36* mutations present in both strains, both strains had a single additional homozygous mutation in common relative to our wild type strain and the reference *C. elegans* sequence. This homozygous mutation was in the autophagy-related gene *atg-9* (*T22H9.2*) that is within the region to which *n3710* mapped on chromosome V.

We carried out a complementation test to determine whether *n3710* is indeed a mutation in *atg-9*. We first crossed wild type worms to *cup-5(zu223) unc-36(e251); n3710* hermaphrodites (*x* = *+* in Fig 1A). Out of 93 F_2_ *cup-5 unc-36* mutant hermaphrodites, 18 (19.4%) laid embryos that hatched and grew, indicating that *n3710* is a recessive mutation. We then crossed *atg-9(bp564)* males to *cup-5(zu223) unc-36(e251); n3710* hermaphrodites (*x* = *atg-9* in Fig 1A). *atg-9(bp564)* is a null allele that results in a Q235 to stop codon change in ATG-9A [49, 50]. 92 (86.7%) of the 106 F_2_ *cup-5 unc-36* mutant hermaphrodites, laid embryos that hatched and grew, indicating that *n3710* does not complement *bp564*. In addition, the *atg-9(bp564)* mutation alone also suppressed the *cup-5(zu223)* embryonic and larval lethality (see Fig 6).

ATG-9 is an autophagy-related gene that is homologous to human Atg9. This family of conserved proteins has six predicted transmembrane domains and functions in autophagy progression from phagophores to autophagosomes, lifespan extension, neuronal modeling, apoptotic cell clearance, and protection from polyQ toxicity [49, 51–54]. All of these functions are thought to be related to ATG-9’s primary function in autophagy. During autophagy, Atg9 is thought to transport lipids from membranes of origin to the phagophore assembly site [55–58].

The *n3710* mutation introduces an A to V change in the fourth predicted transmembrane domain, altering an AAA to an AVA sequence in the ATG-9 protein (Fig 4A). While this change appears conservative, it could affect the lipid transfer capabilities of ATG-9 that likely depend on the association of the transmembrane domains with each other. Indeed, the homozygous null mutation *atg-9(bp564)* does not appear to show a stronger suppression phenotype of *cup-5(null)* lethality, either homozygous or heterozygous with *atg-9(n3710)*, than homozygous *atg-9(n3710)* (see complementation analysis above and Fig 6).

**Fig 4.**
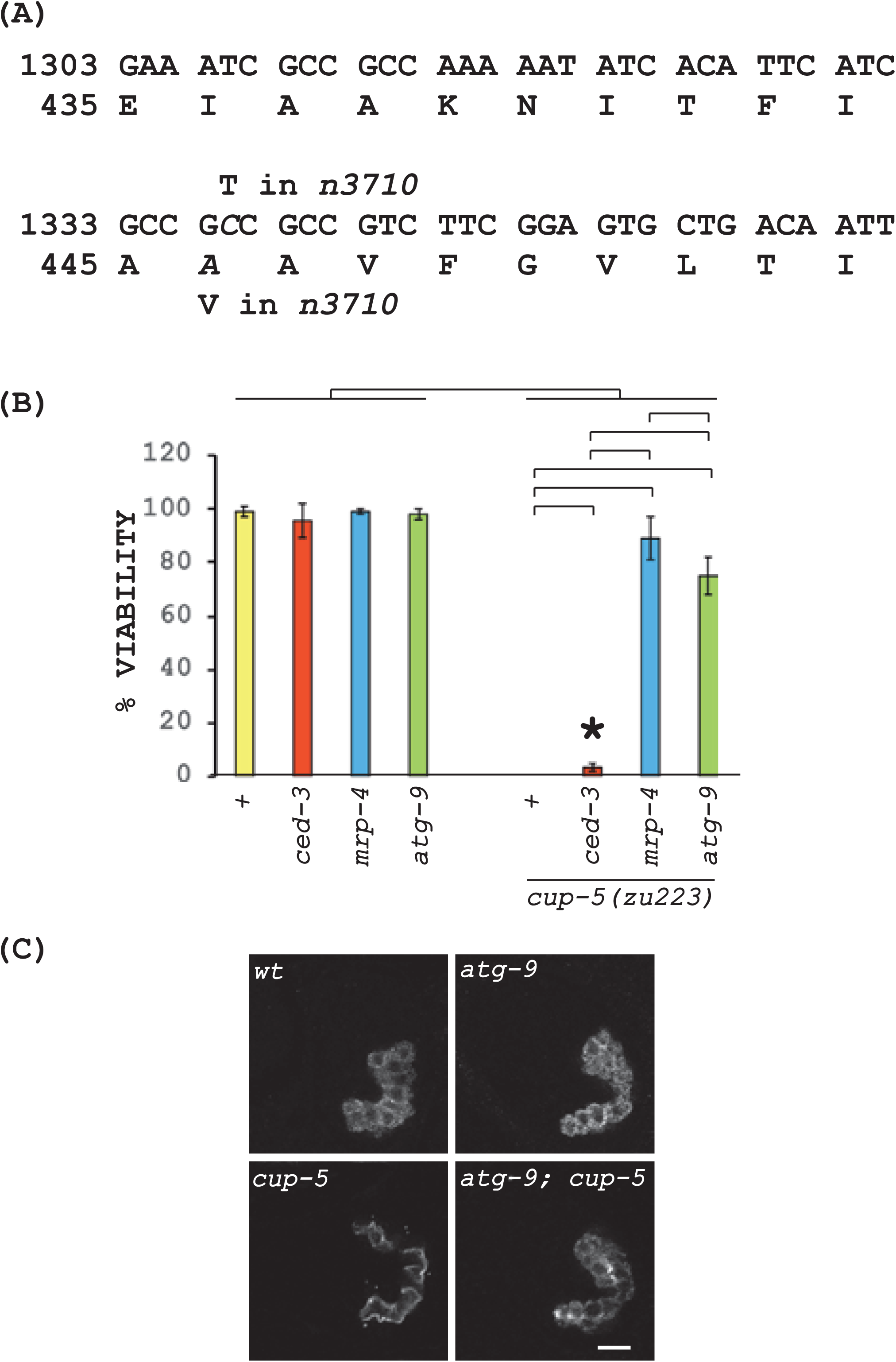
*atg-9(n3710)* sequence change and suppression of *cup-5(null)* cell death. (A) Part of the predicted cDNA sequence of *atg-9* showing the nucleotide and corresponding amino acid changes (italicized) in the predicted protein. (B) Percent of viable worms laid by the indicated wild type, single mutant, and double mutant hermaphrodites. The asterisk indicates that the *ced-3; cup-5* mutant embryos that hatched arrested at the L1 stage. Inverted brackets indicate *P<*0.05; All of the single mutants showed *P<*0.05 relative to all of the double mutant strains. *mrp-4* and *ced-3* mutants are shown for comparison. All strains, except for the *atg-9(n3710)* single mutant, also had the *arIs37* transgene. (C) Confocal images of “comma” to “1.5-fold” stage embryos laid by the indicated hermaphrodites and immuno-stained to detect the intermediate filament protein IFB-2. Scale bar represents 10 µm in whole embryos

*atg-9(n3710)* strongly suppresses the *cup-5(null)* embryonic and larval lethality. While almost 100% of embryos laid by *cup-5(zu223)* single mutants fail to hatch, 75 +/- 7% of *atg-9(n3710); cup-5(zu223)* embryos hatch and develop, albeit at a slightly slower rate than wild type, to fertile adult hermaphrodites (Fig 4B). This is similar to the suppression of *cup-5(zu223)* by *mrp-4(null)* mutants, and contrasts with the weak suppression of *cup-5(null)* by *ced-3(null)* mutant (Fig 4B) [17, 19]. Indeed, *atg-9(n3710)* restores the normal intestinal architecture of *cup-5(null)* worms, as detected by staining for the intermediate filament protein IFB-2. The strong suppression of *cup-5(null)* intestinal cell death and embryonic lethality by *atg-9(n3710*) indicates that normal ATG-9 is necessary for the cell death and embryonic lethality in the absence of CUP-5.

### *atg-9(n3710)* does not suppress the lysosomal defect of *cup-5(null)*

Given the strong suppression of the cell death and lethality of *cup-5(null)*, we expected that the *atg-9(n3710)* mutant would also suppress the lysosomal defect due to the absence of CUP-5. Indeed, loss or reduction of function of MRP-4 or of ESCRT-Associated proteins strongly suppress the cell death/embryonic lethality and the lysosomal defect of *cup-5(null)* (Fig 4B): this indicates that the presence of normal MRP-4 and ESCRT-Associated proteins is necessary for the lysosomal defects in the absence of CUP-5 [19, 20].

We first assayed strains expressing the VIT-2::GFP reporter. VIT-2::GFP is endocytosed by oocytes and subsequently degraded primarily in lysosomes of intestinal cells in developing embryos [39, 59]. In the absence of CUP-5, VIT-2::GFP accumulates in expanded lysosomes that are marked with LMP-1, the worm homologue of Lamp1 [17, 59]. The *atg-9(n3710)* single mutant does not affect the sizes of VIT-2::GFP-positive compartments nor the intensity of VIT-2::GFP in developing intestinal cells relative to wild type, indicating normal membrane transport and lysosomal degradation of VIT-2::GFP in this *atg-9* mutant (Fig 5A, B, C). *cup-5(zu223)*, as expected, showed a significant increase in the sizes of VIT-2::GFP-positive compartments and in the intensity of VIT-2::GFP, indicating defects in lysosome size and lysosomal degradation (Fig 5A, B, C) [17, 59]. Surprisingly, *atg-9(n3710); cup-5(zu223)* embryos showed a size and degradation defect that was indistinguishable from the *cup-5(zu223*) single mutant, indicating a lack of suppression of *cup-5(null)* lysosomal defects (Fig 5A, B, C).

**Fig 5.**
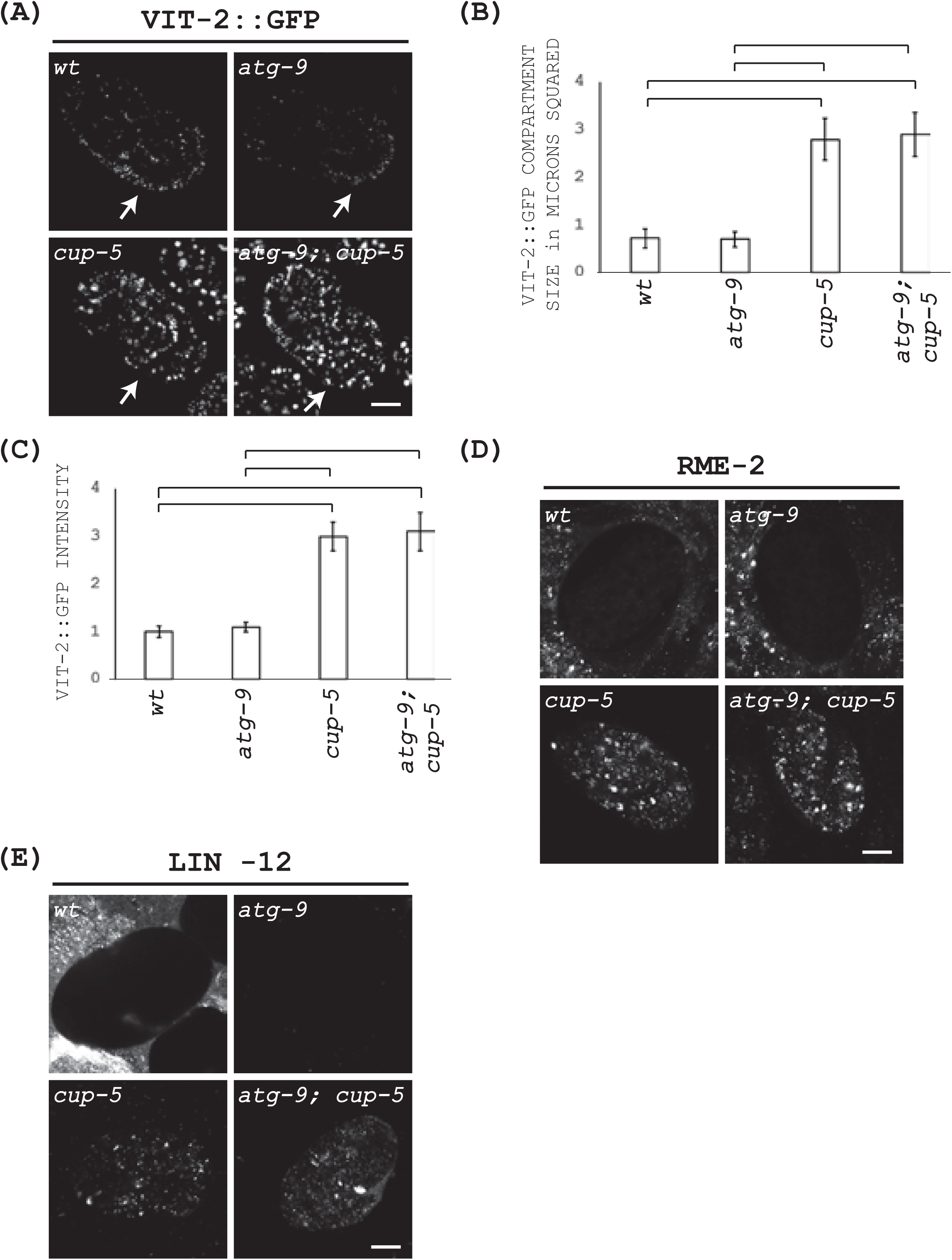
Lysosomal defects in the absence of CUP-5. (A) Confocal images to detect VIT-2::GFP of “comma” to “1.5-fold” stage embryos laid by the indicated hermaphrodites. All images were taken using the same microscopy conditions. The arrow points at the position of the intestinal cells. Scale bar represents 10 µm. (B) Quantitation of the sizes of VIT-2::GFP compartments in developing intestinal cells from panel “A”. (C) Quantitation of the intensities of VIT-2::GFP in developing intestinal cells from panel “A”. The arbitrary intensity units were normalized to “1” using wild type numbers. (D) Confocal images of “comma” to “1.5-fold” stage embryos laid by the indicated hermaphrodites and immuno-stained to detect RME-2. All images were taken using the same microscopy conditions. Scale bar represents 10 µm. (E) Confocal images of “comma” to “1.5-fold” stage embryos laid by the indicated hermaphrodites and immuno-stained to detect LIN-12. All images were taken using the same microscopy conditions. Scale bar represents 10 µm.

To confirm this result, we examined the lysosomal degradation of the yolk receptor RME-2 and the Notch receptor LIN-12 [39, 60]. Indeed, the defective lysosomal degradation of neither RME-2 nor LIN-12 was suppressed by *atg-9(n3710)* in the absence of CUP-5 (Fig 5D, E). These results indicate that in the absence of CUP-5, normal ATG-9 is not required for the lysosomal defects to occur but is required to cause the cell death due to the lysosomal defects.

### *atg-9(n3710)* protects against lysosomal membrane permeabilization

The MSDH results suggest that lysosomal membrane permeabilization is the major reason for cell death in the absence of CUP-5 (Fig 3). Is ATG-9 mediating this lysosomal membrane permeabilization? We first tested cytoplasmic leakage of CPR-6 and GBA-3 using organelle fractionation. Indeed, atg-9(n3710 suppresses the cytoplasmic leakage of both enzymes in the absence of CUP-5 (Fig 2B).

We then tested MSDH-induced lethality, a more functional assay of lysosome membrane permeabilization. *atg-9(n3710)* reduced the effect of MSDH-induced lethality of *cup-5(ar465)* and *cup-5(cd12*) (Fig 3B). Following *cup-5* RNAi, the *cup-5(ar465)* single mutant shows a percent viability of 18 +/- 6 in DMSO and 2.3 +/- 1.2 in MSDH (*P* 0.01); in contrast, following *cup-5* RNAi, the *atg-9(n3710); cup-5(ar465)* double mutant shows a percent viability of 71 +/- 7 in DMSO (showing suppression of *ar465* mutant lethality after *cup-5* RNAi in DMSO) and 63 +/-7 in MSDH (*P* 0.2) (Fig 3B). Following *cup-5* RNAi, the *cup-5(cd12)* single mutant shows a percent viability of 68 +/- 16.7 in DMSO and 4.7 +/- 3 in MSDH (*P* 0.003); in contrast, following *cup-5* RNAi, the *atg-9(n3710); cup-5(cd12)* double mutant shows a percent viability of 91 in DMSO (showing suppression of *cd12* mutant lethality after *cup-5* RNAi in DMSO) and 86.7 +/- 3.5 in MSDH (*P* 0.1) (Fig 3B).

Our results are consistent with loss/reduction of ATG-9 levels suppressing the membrane permeabilization or rupture of defective lysosomes in the absence of CUP-5. Given the known functions of ATG-9 during autophagy, we then asked whether active autophagy is leading to the lysosome membrane permeabilization in the absence of CUP-5, especially since loss of CUP-5 leads to increased autophagy in embryos [17].

### ATG-9 function in the absence of CUP-5 is unrelated to autophagy

If active autophagy is leading to cell death in the absence of CUP-5, then blocking autophagy using other mutations than *atg-9* should also rescue the embryonic lethality of *cup-5(null)*. We tested autophagosome maturation mutants *atg-2(null)*, *atg-3(uncurated deletion)*, *atg-4.1(null)*, *atg-7 (uncurated mutation)*, *atg-9(bp564,* null), and *unc-22(uncurated mutation)* as a control [49, 50, 53, 61–63]. We used a suppression test cross in which significantly more *cup-5(zu223)* F_2_ worms would yield viable progeny than when wild type (x = *+* in Fig 1B) or control worms (*x* = *unc-22* in Fig 1B) are crossed to *cup-5(zu223)* (Fig 1B). Only *atg-9(bp564)* showed suppression of the embryonic viability of *cup-5(zu223*) (Fig 6). This result suggests that ATG-9 is causing lysosomal membrane permeabilization in the absence of CUP-5 in the absence of autophagic flux.

**Fig 6.**
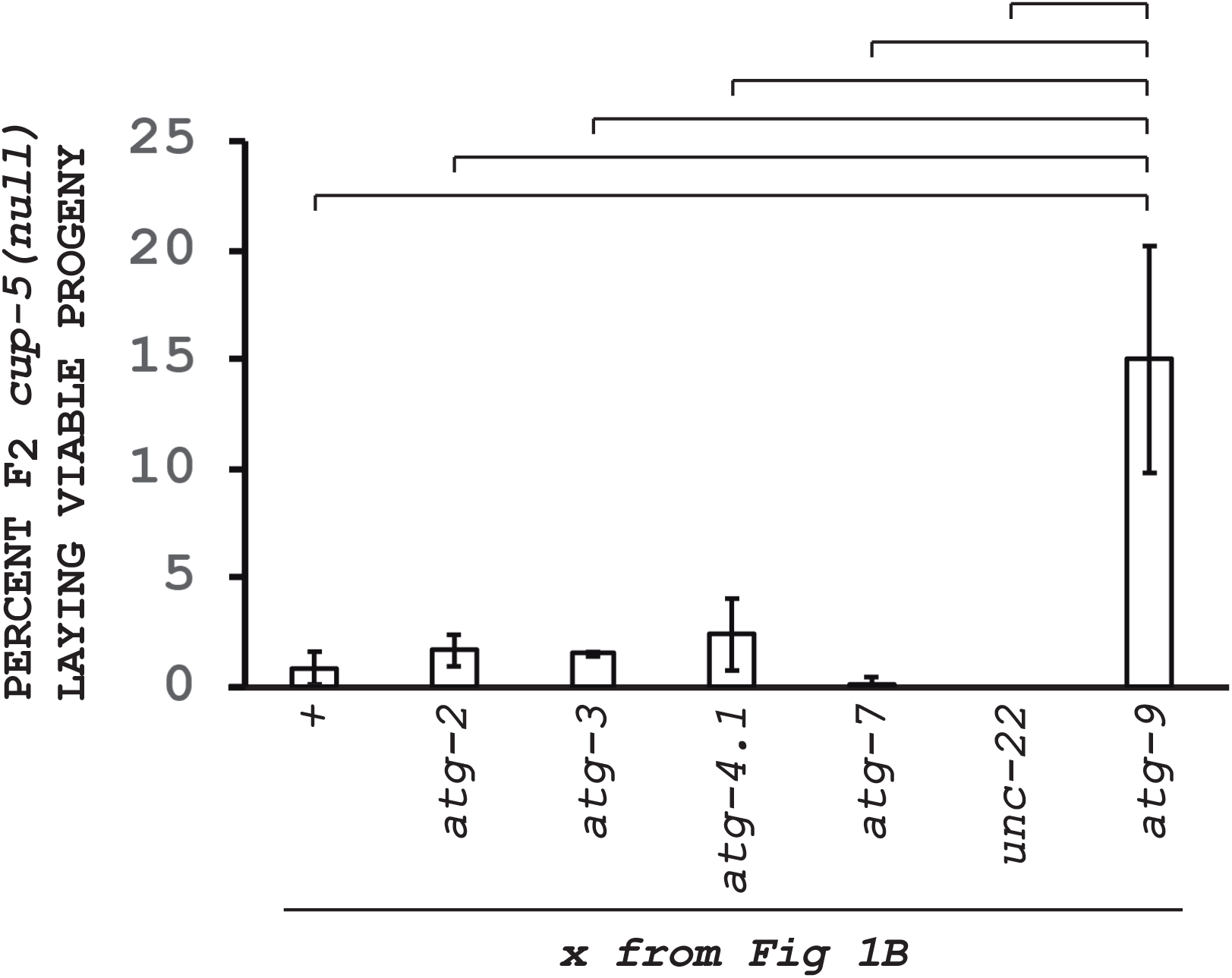
Autophagy mutants suppression of *cup-5(null)* lethality. Percent of *cup-5(zu223)* F2 worms from cross depicted in Fig 1B that laid viable worms for each of the indicated mutations (x in Fig 1B). Inverted brackets indicate *P<*0.05.

How could ATG-9 be mediating this lysosomal permeabilization? There are multiple autophagy defects described in MLIV cells and in worms lacking CUP-5, including constitutive autophagy activation, increased autophagosome formation, and delayed autolysosomal degradation [9, 17]. While these may increase levels of ATG-9 on expanded lysosomes in MLIV patients or *cup-5* mutant worms, our results suggest that autophagy is not required for ATG-9 to exert its lysosomal permeabilization function in the absence of CUP-5. Indeed, mammalian Atg9 is found on late endosomes and the Golgi apparatus under nutrient-rich conditions [64, 65]. This late endosomal localization indicates that it is already present in the right membrane to mediate the permeabilization of the aberrant lysosomes in MLIV or *cup-5* mutant cells. The effect of ATG-9 on lysosomal membranes in the absence of CUP-5 may still be related to its presumed biochemical function in lipid transport between organelle membranes during autophagy [55, 57, 58].

### Model for CUP-5/TRPML1-induced lethality

We generated a model leading from the absence of CUP-5 to cell death (Fig 7A). When CUP-5 is absent, the earliest defect we have detected so far is a hyper-activation of ESCRT-Associated de-ubiquitination leading to the hypo-ubiquitination and thus hyper-activation of the ABC Transporter MRP-4: hyper-activated MRP-4 transports substances into lysosomes causing their dysfunction, and their propensity to permeabilize, in developing intestinal cells (Fig 7A) [19, 20].

**Fig 7.**
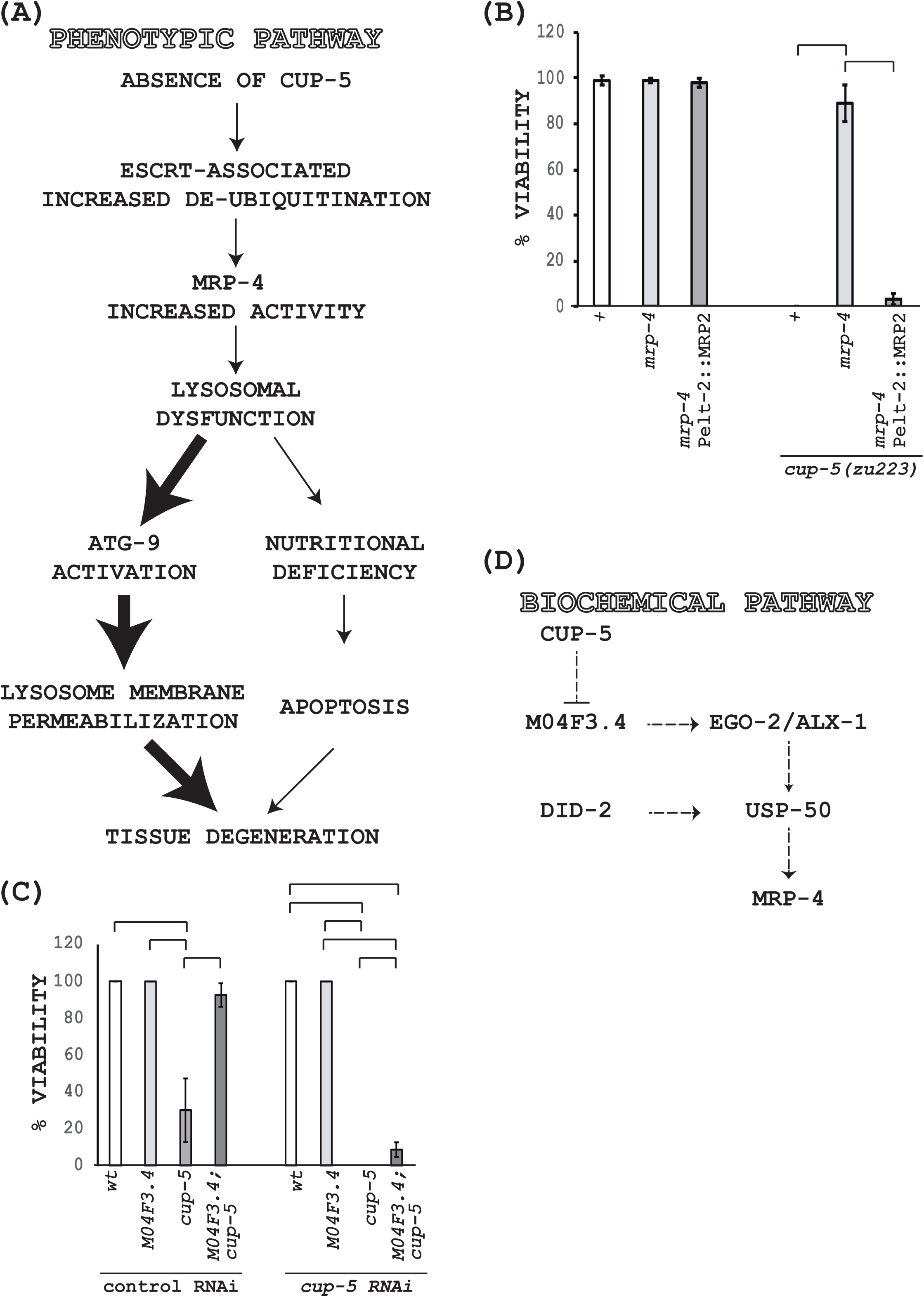
CUP-5 models in developing intestinal cells. (A) Phenotypic/genetic pathway linking CUP-5 to developing intestinal cell death and embryonic lethality. The thicker arrows represent the pathway that more strongly impacts the cell death. (B) Percent of viable worms laid by the indicated hermaphrodites. Inverted brackets indicate *P<*0.05. All strains also had the *arIs37* transgene. (C) Percent of viable worms laid by the indicated wild type and mutant hermaphrodites after control (L4440) or *cup-5* RNAi. Inverted brackets indicate *P<*0.05. All strains also had the *arIs37* transgene. (D) Potential biochemical/physical links between CUP-5 and ESCRT-Associated proteins. Arrows indicate induction; the T-shaped line indicates repression.

To determine whether a human ABC Transporter may show similar functions, we have so far tested the human ABC Transporters MRP2 and it reversed the *mrp-4(null)* suppression of *cup-5(null)* lethality. This indicates that like worm MRP-4, human MRP2 is able to transport the substances into late endosomes/lysosomes that make lysosomes susceptible to rupture/permeabilization (Fig 7B).

Once there is a lysosomal defect, two pathways lead to cell death. First, nutrient starvation activates conventional apoptosis; this pathway accounts for a low percentage of cell death because increasing ATP levels with MethylPyruvate or genetically blocking conventional apoptosis only allows 5-15% of embryos to hatch, but these do not develop beyond the L1 larval stage (Fig 7A) [17]. Second, ATG-9-mediated lysosomal membrane permeabilization is the major reason for the cell death since loss of ATG-9 results in strong suppression of the embryonic and larval lethality (Fig 7A). This pathway, if confirmed in human MLIV patients, may offer some insights into the treatment of MLIV because all of the proteins are conserved in mammals.

Figure 7A shows a genetic model of cell death in the absence of CUP-5. But is there a biochemical link between CUP-5 and the earliest phenotype in the model, the ESCRT-Associated proteins, in normal cells? Human TRPML1 is a calcium channel (among other ions) that associates with the small Ca^2+^-binding protein PDCD6/ALG-2 (Programmed Cell Death-6/Apoptosis Linked Gene-2) [22, 25, 66–69]. We think this TRPML1-ALG-2 connection is relevant for TRPML1^-/-^-mediated cell death because a deletion mutation in *M04F3.4* (encodes the *C. elegans* homologue of ALG-2) suppressed the embryonic and larval lethality of the strong hypomorphic allele *cup-5(bp510)* after control RNAi or *cup-5* RNAi (Fig 7C) [70]. *M04F3.4(null)* does not suppress *cup-5(null)*, suggesting that there is another small Ca^2+^-binding protein in *C. elegans* that has functions relevant to the cell death pathway in the absence of CUP-5. For example, TRPML1 is known to regulate another small Ca^2+^-binding protein calmodulin [71, 72].

Why is this TRPML1-ALG-2 connection relevant? ALG-2 associates with the ESCRT-Associated protein Alix (worm ALX-1) and its homologue HD-PTP (EGO-2) in a Ca^2+^-dependent manner [73–77]. The ALG-2-Alix interaction is thought to be relevant for naturally occurring motor neuron death [78, 79]. *Saccharomyces cerevisiae* studies have shown that Bro1p (homologue of Alix) activates the ESCRT-Associated protein and de-ubiquitinase Doa4p (homologue of mammalian USP8/UBPY and worm USP-50) that is recruited to membranes by the ESCRT-Associated protein Did2p (yeast)/CHMP1 (mammals)/DID-2 (worms) [80–82].

The biochemical model shown in Figure 7D is that in normal cells, CUP-5 inhibits ALG-2 (and another small Ca^2+^-binding protein). ALG-2 normally binds and activates ALX-1/EGO-2 that activate USP-50 that is recruited to endosomal membranes by DID-2. USP-50 de-ubiquitinates MRP-4 thus regulating its activity (our studies suggest that in contrast to normal ESCRT functions, the de-ubiquitination of MRP-4 regulates its activity rather than its levels) [20]. Thus, in the absence of CUP-5, ALG-2 is hyper-active, leading to hyper-activation of the de-ubiquitinase USP-50 and thus the hypo-ubiquitination and over-activation of MRP-4, thus leading to increased transport of substances into late endosomes/lysosomes causing the lysosomal defects (Fig 7A). Reducing the activities of ALX-1/EGO-2, USP-50, DID-2, or MRP-4 strongly rescues the cell death and lethality due to loss of CUP-5 [19, 20].

## Conclusions

Our studies are consistent with a model where the endo-lysosomal defect in the absence of CUP-5 results in the permeabilization, even perhaps rupture, of some lysosomes through an autophagic flux-independent function of ATG-9. This permeabilization is the main reason for the cell death in the absence of CUP-5.

Why do only some tissues die in MLIV patients and in *C. elegans* lacking CUP-5? We think that this is due to a combination of factors. First and foremost is that cell death is restricted to tissues where there is a TRPML1-ALG-2 connection that imposes TRPML1 regulation of the ESCRT-Associated protein Alix (or its homologue HD-PTP). The Alix-ALG-2/PDCD6 connection has already been implicated in neuronal death under normal conditions and in disease states [58, 78, 79, 83–87]. Second, cell death is restricted to tissues that have an ABC Transporter that, when hyper-activated, transports material that make lysosomes susceptible to permeabilization. Indeed, worm MRP-4 expression is restricted to intestinal cells [19]. Worm MRP-4 belongs to the ABCC family of ABC Transporters that includes human MRP1-MRP6 [88, 89]; MRP1-MRP5 mRNA was detected in human brain samples while only MRP1, MRP4, and MRP5 proteins were detectable by immunofluorescence microscopy of brain sections [90]. Our studies suggest that human MRP2 could transport substances that destabilize lysosomes; it remains to be seen whether other ABC Transporters have similar effects on lysosomes when hyper-activated. Third, the presence of Atg9 that causes the permeabilization of lysosomal membrane that have been destabilized by the activity of the ABC Transporter. In mice, Atg9a though ubiquitously expressed, is highly expressed in the central neurons of the nervous system [91]. We think that it is the congruence of these three factors that lead to tissue-specific death in the absence of CUP-5 in worms and in MLIV patients.

Lysosomal membrane permeabilization functions in cell death of normal cells and in disease states [92]. Indeed, lysosomal membrane permeabilization has been implicated in the cell death mechanism of a growing number of lysosomal storage disorders that comprise at least 50 different diseases, including MLIV [92–96]. TRPML1 dysfunction has also been implicated in Niemann Pick diseases, raising the possibility that lysosomal membrane permeabilization in these lysosomal storage diseases could also be mediated by Atg9 [97].

Future studies will uncover Atg9-mediated and Atg9-independent mechanisms of lysosomal membrane permeabilization in lysosomal storage and other diseases.

## Acknowledgments

We thank Brad Hersh and H. Robert Horvitz for the *n3710* suppressor, Susan Miller and Muhammad Noon for whole-genome sequencing and analysis at the University of Arizona, and Gene Dubowchik for the protocol to synthesize MSDH. Several strains were acquired from the Caenorhabditis Genetics Center (CGC). This work was funded by NSF grant MCB-1243037 to H.F.

